# Avian influenza virus surveillance across New Zealand and its subantarctic islands detects H1N9 in migratory shorebirds, but not 2.3.4.4b HPAI H5N1

**DOI:** 10.1101/2024.09.29.615640

**Authors:** Stephanie J Waller, Janelle R Wierenga, Lia Heremia, Jessica A Darnley, Isa de Vries, Jeremy Dubrulle, Zoe Robinson, Allison Miller, Chris N Niebuhr, David Melville, Rob Schuckard, Phil F Battley, Michelle Wille, Rosalind Cole, Kate McInnes, David Winter, Jemma L Geoghegan

## Abstract

Highly pathogenic avian influenza (HPAI) virus has never been detected in New Zealand. The potential impact of this virus on New Zealand’s wild birds would be catastrophic. To expand our knowledge of avian influenza viruses across New Zealand, we sampled wild aquatic birds from New Zealand, its outer islands and its subantarctic territories. Metatranscriptomic analysis of 700 individuals spanning 33 species revealed no detection of HPAI during the annual 2023-2024 migration. A single detection of H1N9 in red knots (*Calidris canutus*) was noted. This study provides a baseline for expanding avian influenza virus monitoring in New Zealand.

## Main text

Since highly pathogenic avian influenza (HPAI) A virus H5N1 first emerged in poultry in China in 1996, global trade of poultry and recent spillovers to wild birds have resulted in the spread of HPAI H5N1 to every European, African and American country (1). Since 2014, the subclade 2.3.4.4 has been the predominant lineage causing epizootic waves resulting in mass mortality events of wild birds and mammals, as well as poultry (1). Oceania remains the only continent yet to detect HPAI H5N1 2.3.4.4.

Until recently, waterfowl have been the avian group believed responsible for long distance movement of avian influenza viruses (including HPAI H5N1 2.3.4.4) (2). However, following the emergence of subclade 2.3.4.4b and associated epidemiological change of this virus, sea- and shorebirds have also contributed to its long distance dispersal (3). New Zealand’s risk of HPAI has previously been considered low due to the absence of migratory waterfowl and its relative geographic isolation (4,5). However, the continued spread of HPAI to new geographic regions, including Antarctica, the increasing role that sea- and shorebirds play in its dispersal, and the rising number of susceptible host species have increased the risk of HPAI being introduced to Oceania.

To date, New Zealand’s avian influenza virus surveillance efforts have largely focused on waterfowl from mainland New Zealand (6), and have not encompassed New Zealand’s offshore and subantarctic islands which host a diverse range of avian species including members of Charadriiformes, Sphenisciformes, Procellariiformes and Pelecaniiformes (3). Although HPAI has never been detected in New Zealand, low pathogenic strains (LPAI) have been frequently detected through this surveillance over the past 20 years (6). Given the global spread of HPAI H5N1 subclade 2.3.4.4b, including to the Antarctic peninsula (7,8), the risk of incursion into New Zealand and its subantarctic islands has increased (3). Due to this risk and the dynamic situation of HPAI H5N1, it is crucial that we expand our knowledge about avian influenza viruses harboured by aquatic birds in New Zealand.

Herein, we aimed to expand New Zealand’s avian influenza virus monitoring by sampling sea- and shorebirds from both mainland New Zealand and its offshore and subantarctic islands. Between November 2023 and March 2024 (i.e. spring to autumn), oropharyngeal and cloacal swabs were collected from 700 individuals across 33 avian species from 13 locations (Figure 1). Following similar protocols to those previously described (9), total RNA was extracted using ZymoBIOMICS MagBead RNA kit (Zymo Research). Equal volumes of extracted RNA were pooled into 207 libraries based on location, avian species and sample type. Pooled RNA was subject to total RNA metatranscriptomic sequencing with ribosomal RNA depletion using Ribo-Zero-Plus. Paired-end 150bp sequencing of the RNA libraries was performed on the Illumina NovaSeqX platform, achieving an average read depth of 62 (range: 41 - 145) million reads per library. Metatranscriptomics has shown to be sensitive for virus detection and characterisation of all viruses infecting animals, including avian influenza viruses (10).

**Figure 1.**
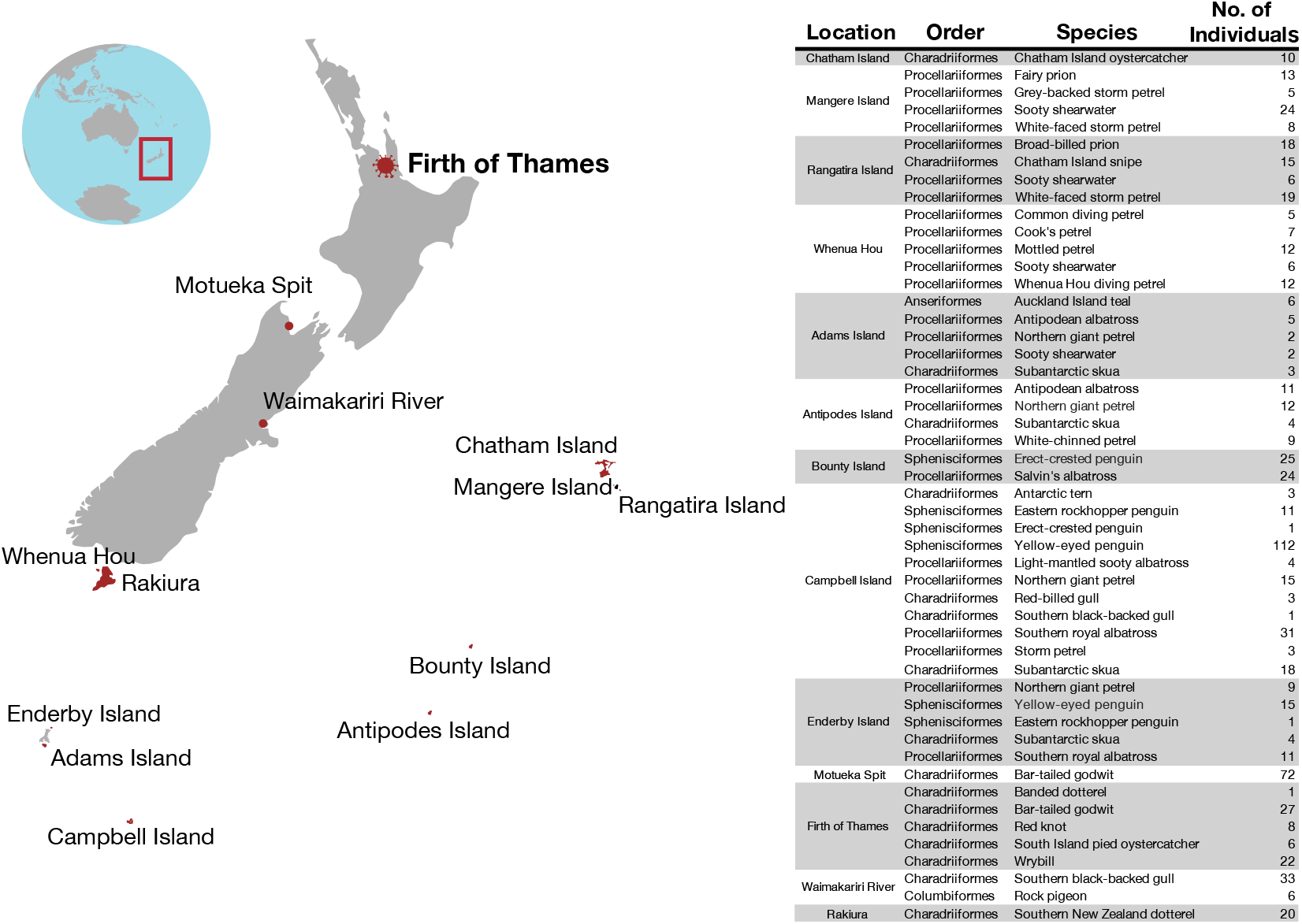
Locations and species sampled during 2023-2024. Map (left) showing the 13 locations where birds were sampled during this study with the Firth of Thames in bold due to the detection of avian influenza virus in birds at this location. Table (right) listing the locations, taxonomic order, species (common name) and number of individual birds that were sampled. Both oral and cloacal swabs were taken from each individual, totalling 1,400 samples.

While HPAI was not detected in any of the 207 pooled libraries via metatranscriptomic sequencing, a near-complete viral genome comprising all eight segments (GenBank accessions: PQ358076-PQ358083) of H1N9 was identified in pooled RNA from cloacal samples from four red knots (*Calidris canutus*) sampled in March 2024 in the Firth of Thames, a site which is part of the East Asian-Australasian Flyway terminus and hosts thousands of migratory birds annually (Figure 2). A time-calibrated maximum likelihood phylogenetic analysis showed that the hemagglutinin (HA) segment fell into a broader Oceania clade with Australian viruses, but diverged from a common ancestor of H1N1 viruses in Australia in the early 1990s. Similarly, the neuraminidase (NA) segment fell into an Oceania clade, related to sequences from both Australia and Asia (Figure 2). Red knots are a migratory species that arrive in New Zealand from mid-September. The detection of this virus in March 2024, combined with the phylogenetic position of HA and NA segments and the relatively large time gap between this virus and the last sampled common ancestors, suggest this LPAI lineage is not a new introduction to New Zealand. However, limited sampling of avian influenza viruses in New Zealand’s migratory birds has hindered our ability to accurately resolve these lineages and determine their likely source. Increasing sampling efforts, particularly throughout the East Asian-Australasian flyway, will help to clarify the evolutionary history and global connectedness of these viruses, enhancing our ability to better understand the transmission dynamics of avian influenza virus throughout this part of the world.

**Figure 2.**
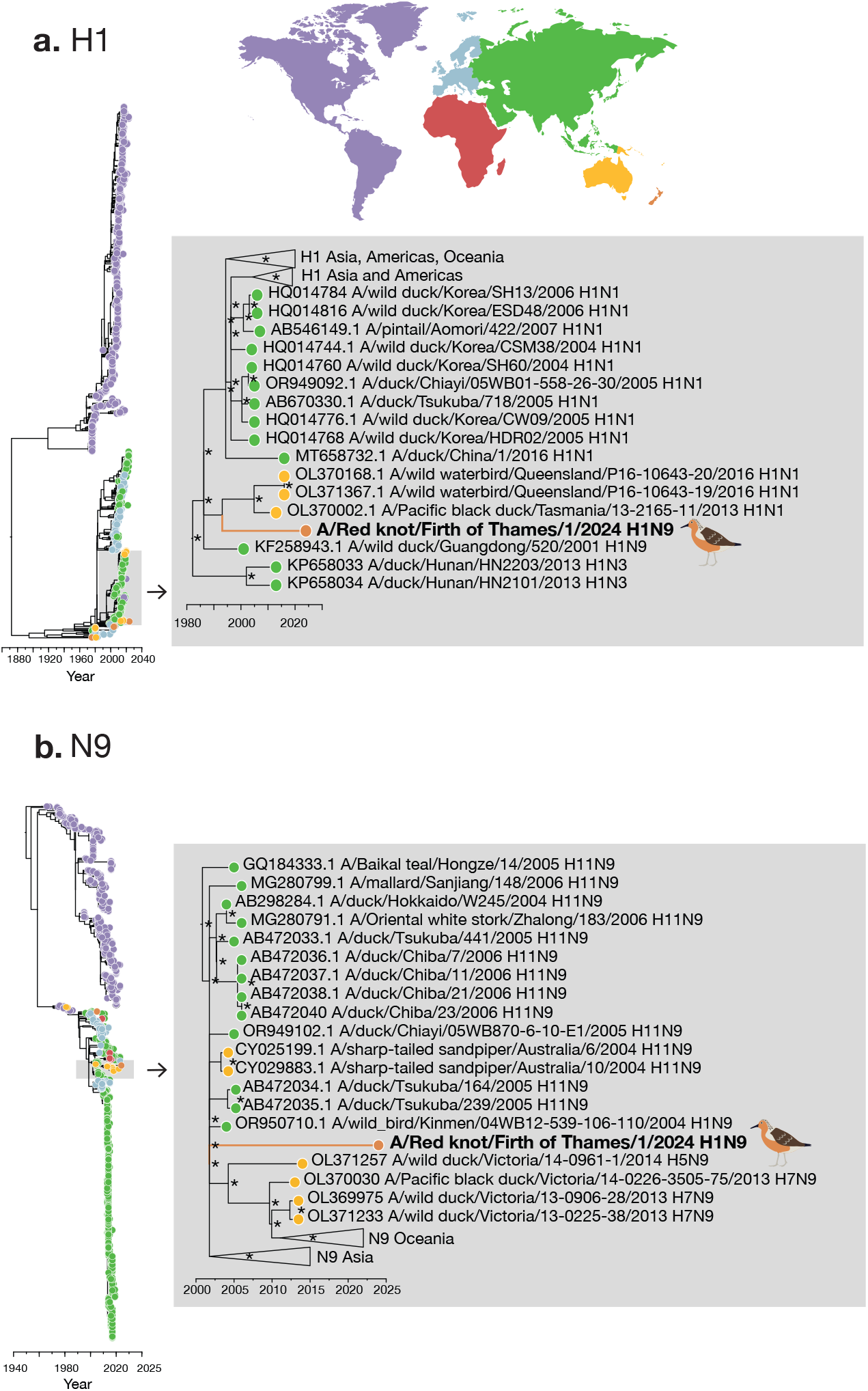
Time-calibrated maximum likelihood phylogenetic trees of H1 and N9 subtypes. Maximum likelihood time-scaled phylogenetic trees using IQ-Tree v1.6.12 (11) of (**a**) H1 and (**b**) N9. Each phylogenetic tree contains the closest 100 genetic relatives from a nucleotide BLAST search as well as all publicly available sequences within each segment subtype (n = 1,332 for H1 and n = 2,010 for N9) obtained from Bacterial and Viral Bioinformatics Resource Center (https://www.bv-brc.org/searches/GenomeSearch), aligned using MAFFT (12). The enlarged images to the right of the phylogenetic trees shows the topological position of Influenza A/Red knot/Firth of Thames/1/2024(H1N9), with ultrafast bootstrapping (13) values of >70%, denoted by an asterisk. Tip colours represent the continents from which the sequences were sampled corresponding to the map above.

The identification of H1N9 in red knots at an internationally important coastal wetland has highlighted the importance of enhanced avian influenza virus surveillance in migratory shorebirds. Approximately 30,000 red knots migrate to New Zealand annually from Siberia, travelling down the East Asian coast, making stopovers in Asia and Australia (14). This migration poses a potential risk for the introduction of HPAI H5N1 into New Zealand. Similar to these findings, a recent investigation in Australia also found high prevalence and seroprevalence of LPAI in red knots as well as ruddy turnstones *(Arenaria interpres*), sharp-tailed sandpipers (*C. acuminat*a) and red-necked stints (*C. ruficollis*), while very low antibody levels were detected in bar-tailed godwits (*Limosa lapponica*), great knots (*C. tenuirostris*), curlew sandpipers (*C. ferruginea*) and sanderlings (*C. alba*) (15). Although these species are from the same avian family, Scolopacidae, the differences in viral prevalence and seroprevalence suggest that there are interspecies differences that may alter avian influenza virus carriage, transmission and exposure potential (15). Nevertheless, with the now wide geographical and host range of HPAI H5N1 subclade 2.3.4.4b, continued surveillance of a broad range of sea- and shorebirds remains critical.

This work has established a framework for the broader surveillance of avian influenza viruses using state of the art genomics approaches, not only in mainland New Zealand, but extending to birds located on New Zealand’s offshore and subantarctic islands. While HPAI was not identified in any of the birds sampled during last season’s 2023-2024 migration to New Zealand, another high-risk period is imminent when migratory birds will arrive back in New Zealand between September – November (5). It is crucial to continue this enhanced surveillance to rapidly detect any viral incursions and build a longitudinal data set to better understand the dynamics of avian influenza virus evolution in New Zealand.

## Author Contributions

**Stephanie J Waller**: Methodology; Investigation; Formal analysis; Visualization; Project administration; Writing – original draft; Writing – review and editing. **Janelle R Wierenga**: Methodology; Investigation; Writing – review and editing. **Lia Heremia**: Methodology. **Jessica A Darnley**: Methodology. **Isa de Vries**: Methodology. **Zoe Robinson**: Methodology. **Jeremy Dubrulle**: Methodology. **Allison Miller**: Writing – review and editing. **Michelle Wille**: Writing – review and editing. **Chris N Niebuhr**: Investigation; Writing – review and editing. **David Melville**: Investigation; Project administration; Resources; Writing – review and editing. **Rob Schuckard**: Investigation; Project administration; Resources; Writing – review and editing. **Phil F Battley**: Investigation; Project administration; Resources; Writing – review and editing. **Rosalind Cole**: Conceptualization; Methodology; Funding acquisition; Project administration; Resources; Writing – review and editing. **Kate McInnes**: Conceptualization; Methodology; Funding acquisition; Project administration; Resources; Writing – review and editing. **David Winter**: Conceptualization; Methodology; Funding acquisition; Project administration; Resources; Writing – review and editing. **Jemma L Geoghegan**: Conceptualization; Methodology; Funding acquisition; Project administration; Resources; Writing – review and editing.

## Acknowledgements

We would like to acknowledge the Department of Conservation, Claudia Mischler, Dave Housten, Graeme Elliott, Jamie Cooper, Johannes Fischer, Kalinka Rexer-Huber, Kath Walker, Kevin Parker, Lydia Uddstrom, Melanie Young, Rachael Sagar, Theo Thompson, Thomas Mattern, Rose Collen, Harrison Talarico, Guy McDonald and Kevin Carter, as well as Environment Canterbury and the crew of Evohe, who were involved in collecting samples for this project.

## Ethics Statement

Animal ethics for sampling birds at the Firth of Thames and Motueka Sandspit were obtained from Massey University Animal Ethics Committee (MUAEC 22/52) and sampling was permitted via the Department of Conservation (Wildlife Act Authority 38111-FAU). All other birds were sampled by the Department of Conservation in accordance with their Wildlife Health Management Standard Operating Procedure.

## References

1. Fusaro A, Zecchin B, Giussani E, Palumbo E, Agüero-García M, Bachofen C, et al. High pathogenic avian influenza A(H5) viruses of clade 2.3.4.4b in Europe - Why trends of virus evolution are more difficult to predict. Virus Evol. 2024;10(1).

2. Yang Q, Wang B, Lemey P, Dong L, Mu T, Wiebe RA, et al. Synchrony of bird migration with global dispersal of avian influenza reveals exposed bird orders. Nat Commun. 2024;15(1):1126.

3. Banyard A, Begeman L, Black J, Breed A, Dewar M, Fijn R, et al. Continued expansion of high pathogenicity avian influenza H5 in wildlife in South America and incursion into the Antarctic region. 2023. Available from: https://www.offlu.org/wp-content/uploads/2023/12/OFFLU-wildlife-statement-no.-II.pdf

4. New Zealand Ministry for Primary Industries. About avian influenza and the risk to New Zealand. Accessed: 2024 Sep 19. Available from: https://www.mpi.govt.nz/biosecurity/pest-and-disease-threats-to-new-zealand/animal-disease-threats-to-new-zealand/high-pathogenicity-avian-influenza/about-avian-influenza-and-the-risk-to-nz/#:∼:text=NewZealand%2CAustralia%2C

5. Williams M, Gummer H, Powlesland R, Robertson H, Taylor G. Migrations and movements of birds to New Zealand and surrounding seas. Wellington: Science & Technical Publishing, Dept. of Conservation; 2006. 1–32 p. Available from: https://www.doc.govt.nz/documents/science-and-technical/sap232.pdf

6. Stanislawek WL, Tana T, Rawdon TG, Cork SC, Chen K, Fatoyinbo H, et al. Avian influenza viruses in New Zealand wild birds, with an emphasis on subtypes H5 and H7: Their distinctive epidemiology and genomic properties. PLoS One. 2024;19(6):e0303756.

7. Fusaro A, Gonzales JL, Kuiken T, Mirinavičiūtė G, Niqueux É, Ståhl K, et al. Avian influenza overview December 2023–March 2024. EFSA J. 2024;22(3).

8. Sub-Antarctic and Antarctic Highly Pathogenic Avian Influenza H5N1 Monitoring Project. Accessed: 2024 Sep 20. Available from: https://scar.org/library-data/avian-flu

9. Waller SJ, Butcher RG, Lim L, McInnes K, Holmes EC, Geoghegan JL. The radiation of New Zealand’s skinks and geckos is associated with distinct viromes. BMC Ecol Evol. 2024;24(1):81.

10. Wille M, Eden JS, Shi M, Klaassen M, Hurt AC, Holmes EC. Virus-virus interactions and host ecology are associated with RNA virome structure in wild birds. Mol Ecol. 2018; 27(24):5263–5278.

11. Nguyen L-T, Schmidt HA, von Haeseler A, Minh BQ. IQ-TREE: A fast and effective stochastic algorithm for estimating maximum-likelihood phylogenies. Mol Biol Evol. 2015;32(1):268–74.

12. Katoh K, Misawa K, Kuma K, Miyata T. MAFFT: a novel method for rapid multiple sequence alignment based on fast Fourier transform. Nucleic Acids Res. 2002;30(14):3059–66.

13. Minh BQ, Nguyen MAT, von Haeseler A. Ultrafast Approximation for Phylogenetic Bootstrap. Mol Biol Evol. 2013;30(5):1188–95.

14. Riegen AC, Sagar PM. Distribution and numbers of waders in New Zealand, 2005–2019. Notornis, 2020;67:591–634.

15. Wille M, Lisovski S, Roshier D, Ferenczi M, Hoye BJ, Leen T, et al. Strong host phylogenetic and ecological effects on host competency for avian influenza in Australian wild birds. Proc R Soc B Biol Sci. 2023;290(1991).

